# Online synaptic credit assignment in active dendrites

**DOI:** 10.64898/2026.09.14.751357

**Authors:** Gan He, Shenjian Zhang, Kai Du, Tiejun Huang

## Abstract

Credit assignment in neural networks is usually formulated as the computation of an abstract error gradient. Whether such a gradient can take a physical, causal form in biophysically detailed multi-compartment neuron models, and enable online, supervised learning, remains unclear. Here we show that the gradient of a detailed neuron’s voltage with respect to a synaptic weight is itself a voltage. Differentiating the discrete backward-Euler update solved by standard simulators yields equations of the same form as the original voltage dynamics, driven by gradient currents. Replaying weight-specific gradient currents forward in time reproduces the exact gradient with high fidelity in an L5 pyramidal neuron across diverse input regimes (*R*^2^ *>* 0.998). Pairing the replayed gradient voltage with a local learning signal yields a causal, online learning rule. A single L5 pyramidal neuron with active dendrites learns to reproduce target voltage trajectories containing calcium plateaus and bursts, and recurrent networks of detailed neurons learn to generate target temporal patterns, far outperforming a readout-only control. These results demonstrate that synaptic credit assignment can be implemented online by the voltage dynamics of detailed neurons, potentially suggesting a physical substrate for gradient-based supervised learning in the brain.

## 1 Introduction

Learning in neural networks requires credit assignment: when the output deviates from a target, each synapse must be assigned its share of the error [Schmidhuber, 2015, Minsky, 1961]. In artificial networks, this is achieved by backpropagation [Rumelhart et al., 1986], which propagates the error backward as a gradient, an abstract quantity that describes how each weight should change. Whether such signals can take a physical, causal form in the brain, particularly in the biophysically detailed neurons used to model it, remains unclear [Lillicrap et al., 2020].

Detailed, multi-compartment neuron models reproduce the active, compartmentalized computations of real dendrites, including calcium plateaus and burst firing [Larkum et al., 2009, Hay et al., 2011]. However, their input-output function is strongly nonlinear and time-varying, making gradients difficult to obtain [Bicknell and Häusser, 2021]. Existing approaches trade accuracy against cost: simplified methods give cheap but approximate gradients [Moldwin and Segev, 2020, Zhang et al., 2023], whereas automatic differentiation give exact gradients but are expensive and offline [Bicknell and Häusser, 2021, Deistler et al., 2025]. Differentiable simulators such as Jaxley [Deistler et al., 2025] address this by running the implicit-Euler solver in an automatic-differentiation framework, so that the gradient of the tridiagonal voltage solve is obtained by backpropagation. This computes exact gradients for arbitrary ion-channel, synaptic, and morphological parameters, and has enabled gradient descent on detailed and network-scale models. However, backpropagation is an offline computation: it stores and then reverses the computation graph, and its large-scale recurrent-network training has relied on simplified dendritic morphologies. A causal, online route from voltage dynamics to synaptic credit assignment, one that does not unroll the computation graph, has been missing.

Here we show that the gradient of a detailed neuron’s voltage with respect to a synaptic weight is itself a voltage. Differentiating the discrete backward-Euler update solved by standard simulators [Hines and Carnevale, 1997] yields equations of the same form as the original voltage dynamics, driven by gradient currents. Replaying these currents forward in time returns the synaptic gradient exactly. Pairing the replayed gradient voltage with a local learning signal produces an online, causal learning rule, enabling training of a single L5 pyramidal neuron with active dendrites and recurrent networks of detailed neurons to generate target dynamics.

## 2 Results

### 2.1 Forward gradient replay reproduces ground-truth gradients across input regimes

Gradient computation in detailed, multi-compartment neuron models proves difficult because the sensitivity of a somatic voltage to a synaptic weight must be propagated through strongly nonlinear, time-varying conductances. We reasoned that this abstract sensitivity might nevertheless have a concrete physical counterpart. In the discrete update equations solved by standard simulators, the voltage at each time step is obtained by solving a system of equations [Hines, 1984]. Its structure is fixed by the morphology and by conductances evaluated at the previous state. Differentiating this system with respect to any synaptic or ionic weight *w*_*i*_ yields a system of the same form as the original, driven by gradient currents *Î*_*j*|*i*_. Substituting the derivative of the voltage with a new variable, the gradient voltage 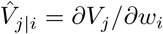, we find that the gradient dynamics is governed by the same equation as the voltage dynamics (Eq. 1). The only difference is that the driving currents are replaced by gradient currents. Because gradient currents superpose, the equivalence holds regardless of how many ionic and synaptic conductances a compartment carries, and it applies equally to ionic-channel and synaptic weights.

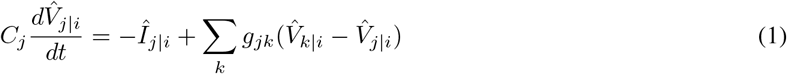

We refer to the resulting procedure as forward gradient replay (Fig. 1a). The voltage simulation is run once, and the state variables of every current source (gating variables, conductances, and related quantities) are recorded simultaneously. These recorded dynamics are used to compute how the state variables of each source evolve during the replay, and the resulting changes generate the gradient currents that drive the gradient voltage. When the gradient is computed with respect to a given synaptic weight, the target synapse and all other current sources replay their gradient currents. The voltage read out from the replay is therefore the gradient of the original voltage with respect to the weight under consideration.

**Figure 1.**
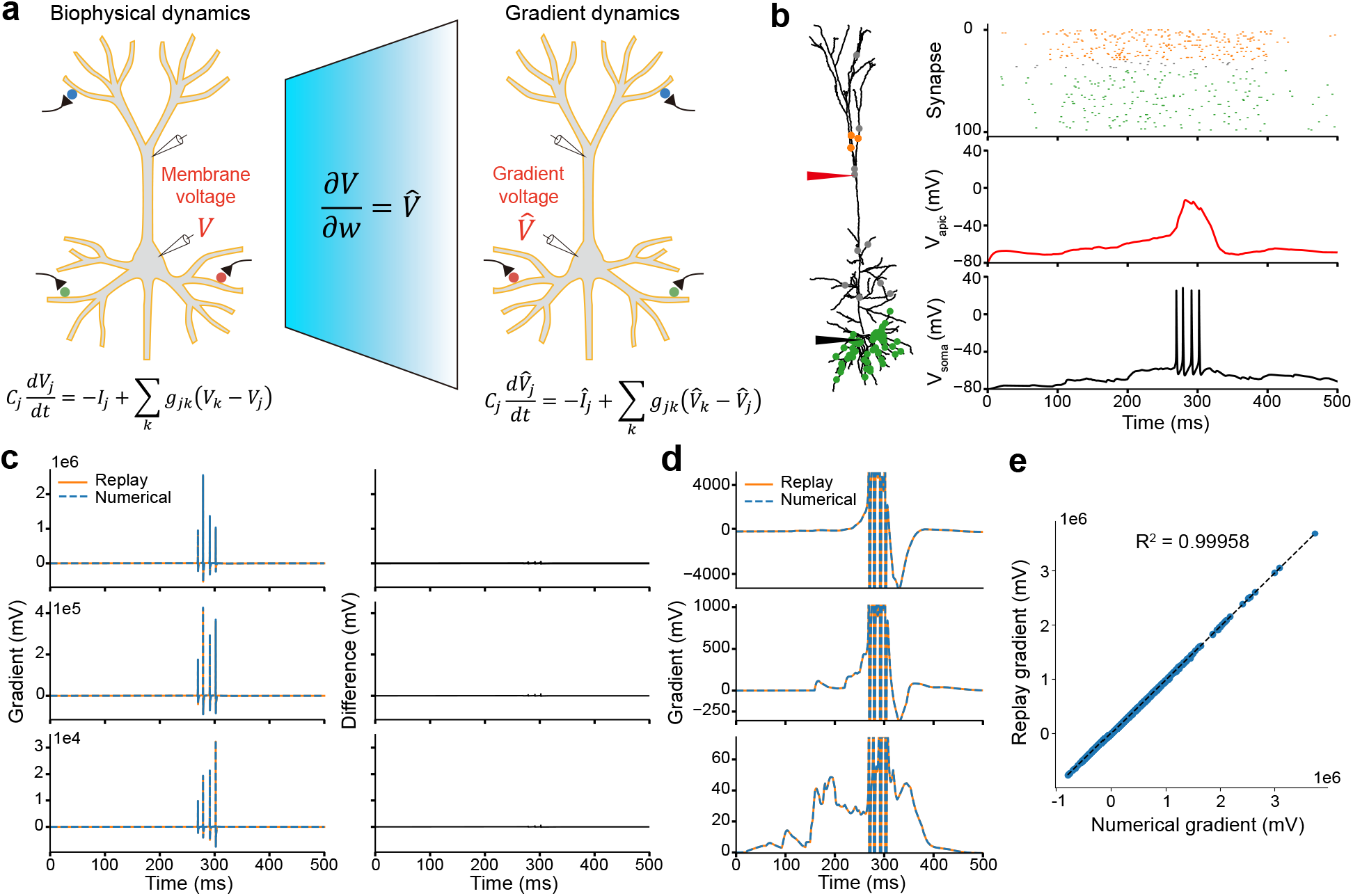
Numerical validation of forward gradient replay. (a) Forward gradient replay scheme. (b)-(e) Validation in high-frequency input regime. (b) Synaptic locations, input spikes and somatic/apical voltage recordings. (c) Gradient traces overlapped. (d) Subthreshold view of gradient traces overlapped. (e) *R*^2^ compared between forward-replay and numerical gradients.

To test this procedure, we used a biophysically detailed rat L5 pyramidal neuron model with active conductances distributed across the soma, apical, and basal dendrites [Hay et al., 2011]. We tested whether forward gradient replay returned gradients that matched ground-truth gradients obtained by numerical differentiation. Three input regimes were considered: random, synchronous, and high-frequency (Fig. 1b, Supplementary Fig. 1a,e). In the random regime, synapses were uniformly distributed; in the synchronous regime, a synchronized spike arrived at clustered apical inputs; and in the high-frequency regime, clustered apical inputs received high-frequency Poisson input. For the synchronous and high-frequency regimes, the neuron received a mix of apical-cluster, apical-background, and basal synaptic inputs, producing an apical calcium plateau and a somatic burst, a hallmark of active dendritic integration. Across all nine representative weights spanning all three regimes, the forward-replay and numerical gradients overlapped throughout the simulation, and the only visible deviations occurred at somatic spikes (Fig. 1c, Supplementary Fig. 1b,f). We then isolated the subthreshold component by retaining the central 95% of each gradient’s data points; within this component the forward-replay and numerical gradients were nearly indistinguishable (Fig. 1d, Supplementary Fig. 1c,g). Across all synaptic weights and all three regimes, the scatter of forward-replay gradients against numerical gradients yielded *R*^2^ *>* 0.998 (Fig. 1e, Supplementary Fig. 1d,h).

### 2.2 Forward gradient replay implements online credit assignment in a single neuron

Forward gradient replay provides, at every time step, the sensitivity of the somatic voltage to each synaptic weight. To turn this into online supervised learning, we paired the replay with a local learning signal. At each time step *t*, the output and target voltages are fed into an online learning-signal generator that produces a neuronal-specific guidance signal 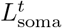. The update of each weight is obtained from this signal together with its gradient voltage. For a neuron with *N* synapses, *N* gradient neurons run in parallel. Each receives the gradient-current state variables transferred from the inference simulation and, at time *t*, the same guidance signal 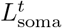. An instantaneous update 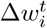 is therefore produced for every synapse (Fig. 2a, Eq. 2). Because the gradient currents carry the neuron’s history forward in time, the update is computed causally, step by step, without unrolling the computation graph.

**Figure 2.**
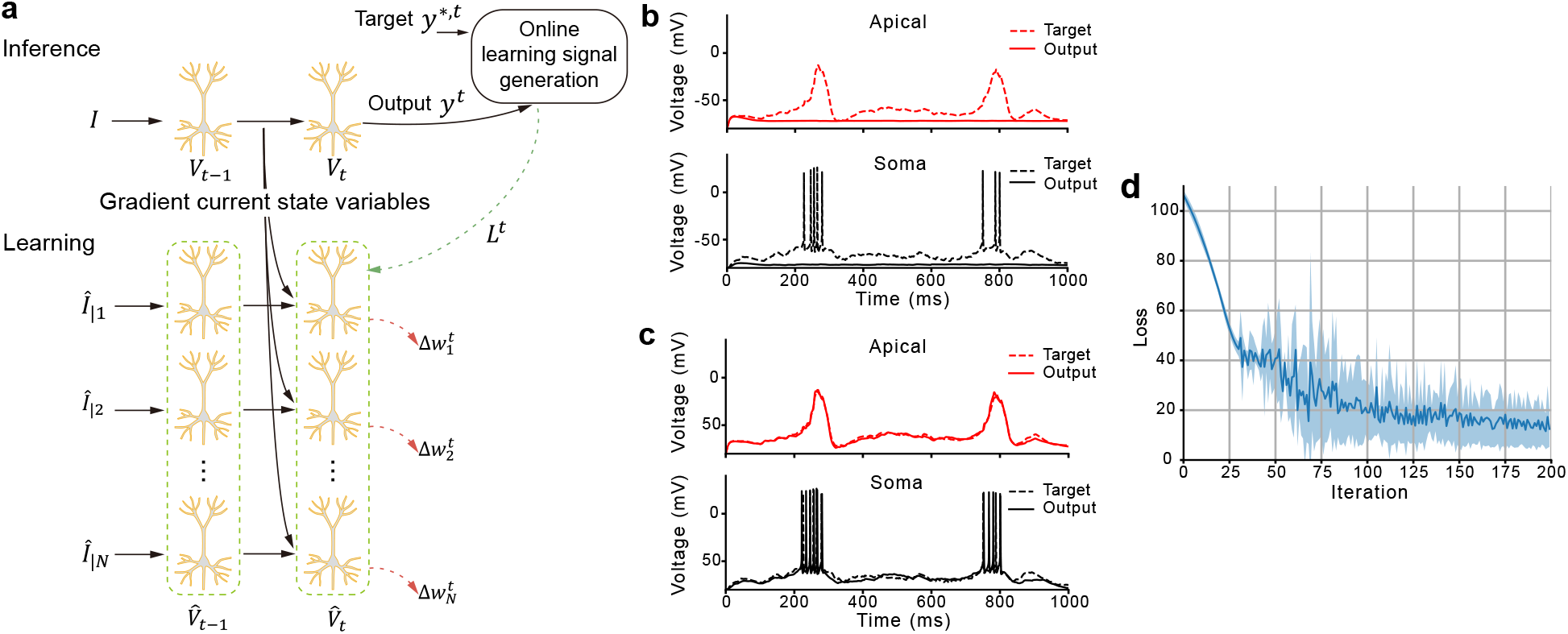
Forward gradient replay enables online credit assignment at single-neuron level. (a) Single-neuron-level weight update scheme. (b) Membrane voltages before learning. (c) Membrane voltages after learning. (d) Loss curve during learning (mean and standard deviation across 5 initializations).

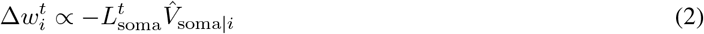

We validated this scheme by training the same rat L5 pyramidal neuron to reproduce target somatic and apical-tuft voltage traces. The target was generated in a high-frequency input regime extended so that the apical dendrite produced two calcium plateaus with two accompanying somatic bursts. Starting from random synaptic weights, the trained neuron closely reproduced the target dendritic-integration function (Fig. 2b,c), and the loss decreased over 200 iterations (Fig. 2d). Forward gradient replay therefore turned a detailed neuron with active dendrites into a trainable unit.

### 2.3 Forward gradient replay trains recurrent networks of detailed neurons online

To test whether the scheme scales to networks, we used a pattern-generation task. A recurrent network of detailed neurons received clockwise Poisson spike inputs and drove a linear readout. The readout produced a target waveform formed by a normalized sum of sinusoids (Fig. 3a). We built two versions of this network, one from mouse L2/3 pyramidal neurons [Bicknell and Häusser, 2021] and one from rat L5 pyramidal neurons. Input, recurrent and output weights were all trained online, with a firing-rate regularization term on the network neurons; a control condition trained only the readout weights. Both networks converged: the L2/3 network converged more smoothly and achieved a closer fit (Fig. 3b,c), and the loss decreased over 300 iterations to values far below those of the readout-only control, which remained high throughout training (Fig. 3d,e). The results demonstrate that the dendritic weights themselves, not merely the readout, contributed to the solution.

**Figure 3.**
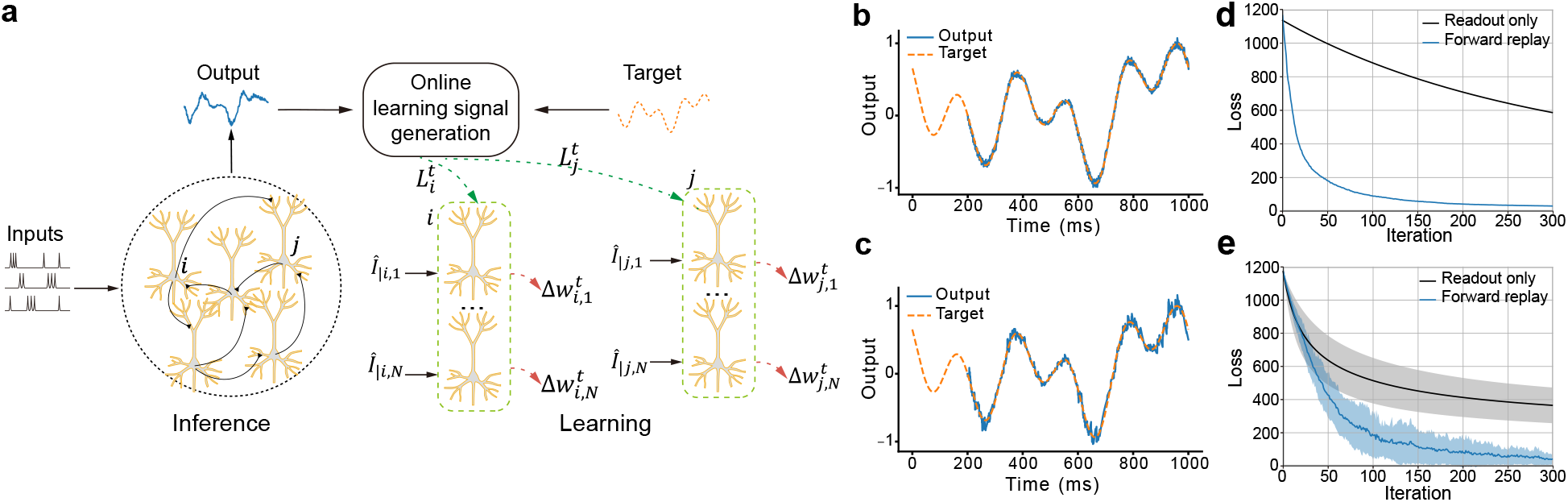
Forward gradient replay enables online credit assignment in detailed recurrent networks. (a) Network-level weight update scheme. (b) L2/3 network generated patterns after training. (c) L5 network generated patterns after training. (d) L2/3 network loss curve during training. (e) L5 network loss curve during training (mean and standard deviation across 5 initializations).

## 3 Discussion

Credit assignment is usually discussed as a computational problem, in which an error is transformed into changes of synaptic weights. Our results show that it can also be discussed as a physical problem, in which the credit signal is carried by the very quantities a neuron already manipulates. We found that the gradient of a voltage with respect to a synaptic weight is itself a voltage, driven by a gradient current that can be replayed forward in time. This renders the gradient a state of the neuron itself, with a defined location, time course and effect on subsequent dynamics. It turns gradient computation from an offline numerical operation into a causal, online process. Because the gradient is a voltage, it is in principle measurable, perturbable and available to downstream computation within biological constraints.

The contrast with differentiable simulators is instructive. Simulators such as Jaxley compute the same gradients by automatic differentiation, but do so by backpropagation, which stores and then reverses the computation graph. This is exact and scalable in parameter count, but it is offline and non-causal, and its large-scale recurrent-network training has relied on simplified dendritic morphologies. Forward gradient replay, by contrast, carries the gradient forward in time together with the neuron’s own state, so that weight updates are produced causally and online in full morphologically detailed, active-dendrite networks. The two approaches are complementary: backpropagation is a general-purpose engine for fitting models, whereas forward gradient replay is a candidate physical mechanism by which a neuron could compute its own credit signals.

This interpretation resonates with the activity-dependent plasticity observed in the brain. Forward replay, in which a sequence of activity is replayed in its original temporal order, has been recorded in the hippocampus during rest and sleep and is thought to support memory and learning [Skaggs and McNaughton, 1996, Ólafsdóttir et al., 2018]. Our framework provides a potential computational explanation for such activity: replaying gradient currents forward in time is how a neuron could compute the credit of its synapses. The learning rule also inherits a well-known approximation. In networks we retain only the direct influence of a weight on its own neuron’s output, in the same spirit as e-prop [Bellec et al., 2020] and related local learning rules.

Several limitations temper these conclusions. First, all results come from simulations. Second, the learning signal that drives each weight update is generated by comparing the neuron or network output with a target, an element that is not fully local. Third, the cost of forward gradient replay scales with the number of synapses, because each weight requires its own gradient simulation. Reducing this cost while preserving the online, causal character of the method is an open direction. Finally, the tasks considered, membrane-potential fitting and pattern generation, are tractable but restricted, and whether the same mechanisms scale to richer tasks remains open.

Together, these results suggest the voltage dynamics of detailed neurons can potentially act as a physical substrate for online synaptic credit assignment. They suggest that abstract gradients need not be imposed on the brain from outside: they can be replayed as voltages by the neurons themselves.

## 4 Methods

### 4.1 Neuron models and synaptic inputs

Simulations used two morphologically detailed neuron models. The rat L5 pyramidal neuron model [Hay et al., 2011] carried active conductances in the soma, apical dendrites and basal dendrites. These were sufficient to reproduce NMDA spikes, calcium plateau potentials, backpropagating action potentials and somatic bursting induced by coincident apical and basal input. The mouse L2/3 pyramidal neuron model [Bicknell and Häusser, 2021], with active conductances confined to the soma, provided a simpler comparison. Excitatory synapses comprised AMPA and NMDA components with reversal potentials of 0 mV and an NMDA:AMPA weight ratio of 2:1; inhibitory synapses were GABAergic with an inhibitory reversal potential. Synaptic and channel parameters followed the published detailed-network and channel models cited above.

### 4.2 Voltage simulation and forward gradient replay

Voltage dynamics were simulated with the backward-Euler implicit scheme used by standard simulators [Hines and Carnevale, 1997]. In this scheme, the voltage at each time step is obtained by solving a system of equations determined by the cable properties and by conductances evaluated at the previous state. Differentiating this system with respect to a synaptic or ionic weight yields equations of the same form as the voltage update. The derivative of the voltage is substituted with a gradient voltage 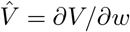, and the resulting equations are driven by gradient currents *Î*.

To compute a gradient in practice, the forward simulation was run once and the state variables of all current sources (gating variables, conductances and related quantities) were recorded. These recorded dynamics were used to compute the gradient currents of the corresponding sources, which were replayed forward in time through a morphologically identical gradient neuron. The gradient neuron had the same compartment geometry, membrane capacitance and axial resistance, with the leak reversal potential set to zero, and the resulting gradient voltage was read out at the desired location. Gradient computation for all weights of a single neuron was parallelized across multiple gradient neurons using MPI.

Gradient accuracy was assessed against numerical differentiation, in which each weight was perturbed by a small amount and the gradient estimated by finite differences. Across all input regimes, the coefficient of determination between forward gradient replay and numerical gradients exceeded 0.999.

### 4.3 Learning rules

At each time step, a guidance signal *L*^*t*^ was generated online from the output and target of the neuron or network. Each weight received an update equal to the product of *L*^*t*^ and its gradient voltage, accumulated over time. In networks, only the direct gradient terms were retained, the influence of a weight on its own neuron’s output, following the approximation used in e-prop [Bellec et al., 2020]. Losses comprised the mean squared error between output and target, with spike-associated values clamped in the forward-replay setting, plus a firing-rate regularization term with a target rate of 10 Hz in network tasks. Weights were updated by stochastic gradient descent with task-specific learning rates; forward-replay gradients were clipped to remove spike-associated extremes.

### 4.4 Tasks

#### Numerical accuracy

Forward gradient replay was compared with numerical differentiation under three input regimes each lasting 500 ms. In the random regime, synapses were uniformly distributed over basal and apical dendrites, receiving 10 Hz Poisson inputs. In the synchronous regime, a synchronized spike was delivered to clustered apical inputs. In the high-frequency regime, clustered apical inputs received 30 Hz Poisson input. Basal and background inputs were low-frequency Poisson in synchronous and high-frequency regimes, being 10 Hz and 2 Hz, respectively. Subthreshold gradients were isolated by retaining the central 95% of each gradient’s data points.

#### Membrane-potential fitting

A rat L5 pyramidal neuron received 30 Hz high-frequency apical-cluster input alongside 10 Hz basal input and 2 Hz background input, with an extended simulation time of 1000 ms, arranged so that the target response contained two apical calcium plateaus and two accompanying somatic bursts. Synaptic weights were initialized randomly and trained for 200 iterations to minimize the mean squared error between the simulated and target somatic and apical-tuft voltages. Somatic spikes were truncated at −40 mV. Results were averaged over five random initializations.

#### Pattern generation

One hundred spike inputs (2 Hz background Poisson noise with 100 ms, 80 Hz high-frequency clockwise bursts whose onsets uniformly covered 100–1000 ms) projected to a recurrent network of 100 detailed neurons. The somatic voltages were read out through a single linear unit with a 30 ms time constant to produce a target waveform formed by a normalized sum of sinusoids. Recurrent connections were made with 10% probability and no self-connections; input and readout connections were all-to-all, with one synapse per connection and a 4:1 excitatory:inhibitory ratio. Networks built from mouse L2/3 or rat L5 pyramidal neurons were each trained for 300 iterations, with a readout-only control condition; L5 results were averaged over five random initializations.

## Supplementary material

**Supplementary Figure 1.**
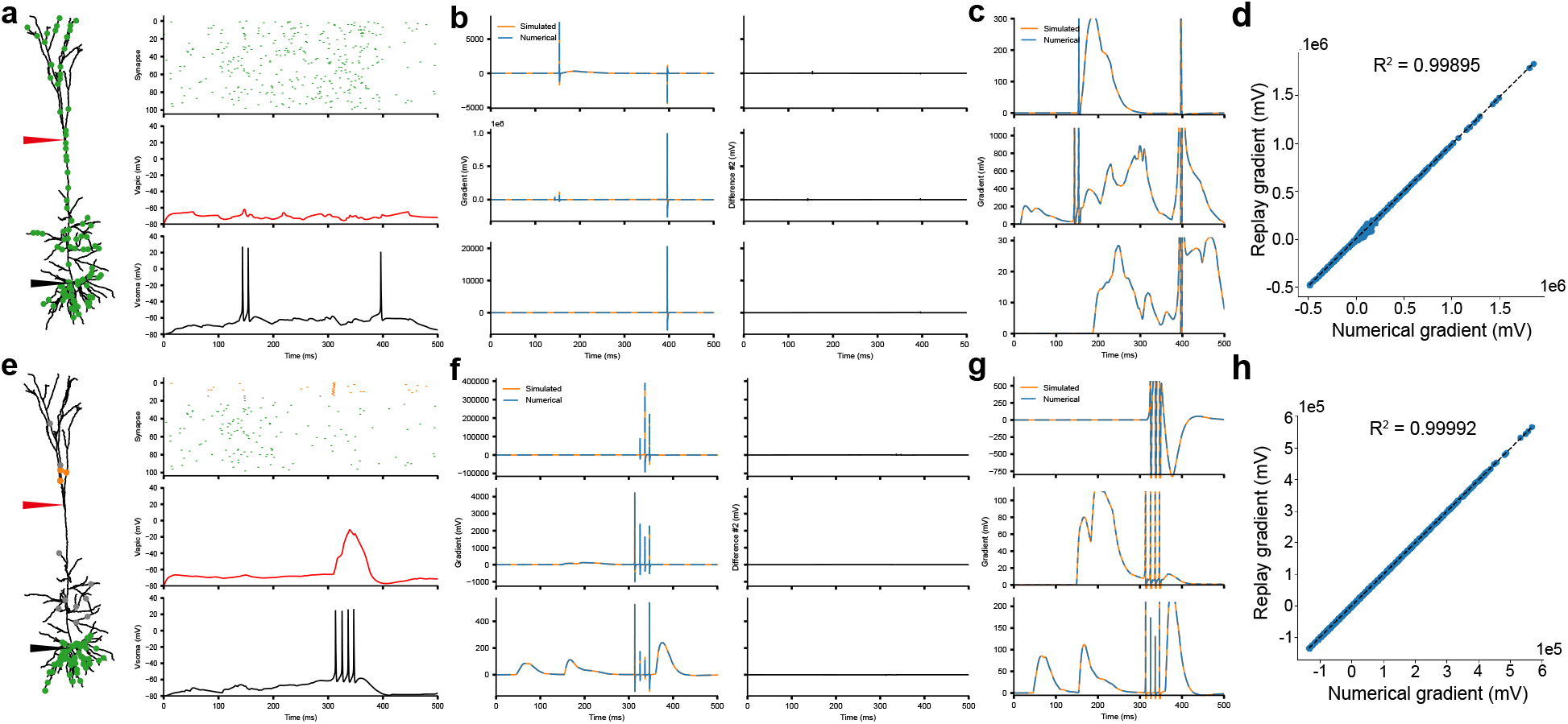
Additional numerical validation of forward gradient replay. (a)-(d) Random input regime. (e)-(h) Synchronous input regime. (a) Synaptic locations, input spikes and somatic/apical voltage recordings. (b) Gradient traces overlapped. (c) Subthreshold view of gradient traces overlapped. (d) *R*^2^ compared between forward-replay and numerical gradients. (e)-(h) Same as (a)-(d) but for synchronous input regime.

